# HIV-1 Gag colocalizes with euchromatin histone marks at the nuclear periphery

**DOI:** 10.1101/2023.02.24.529990

**Authors:** Jordan Chang, Leslie J. Parent

## Abstract

The retroviral Gag protein of human immunodeficiency virus type 1 (HIV-1) plays a central role in the selection of unspliced viral genomic RNA for packaging into new virions. Previously, we demonstrated that full-length HIV-1 Gag undergoes nuclear trafficking where it associates with unspliced viral RNA (vRNA) at transcription sites. To further explore the kinetics of HIV-1 Gag nuclear localization, we used biochemical and imaging techniques to examine the timing of HIV-1 entry into the nucleus. We also aimed to determine more precisely Gag’s subnuclear distribution to test the hypothesis that Gag would be associated with euchromatin, the transcriptionally active region of the nucleus. We observed that HIV-1 Gag localized to the nucleus shortly after its synthesis in the cytoplasm, suggesting that nuclear trafficking was not strictly concentration-dependent. Furthermore, we found that HIV-1 Gag preferentially localized to the transcriptionally active euchromatin fraction compared to the heterochromatin-rich region in a latently-infected CD4+ T cell line (J-Lat 10.6) treated with latency-reversal agents. Interestingly, HIV-1 Gag was more closely associated with transcriptionally-active histone markers near the nuclear periphery, where the HIV-1 provirus was previously shown to integrate. Although the precise function of Gag’s association with histones in transcriptionally-active chromatin remains uncertain, together with previous reports, this finding is consistent with a potential role for euchromatin-associated Gag molecules to select newly transcribed unspliced vRNA during the initial stage of virion assembly.

**Importance:** The traditional view of retroviral assembly posits that HIV-1 Gag selection of unspliced vRNA begins in the cytoplasm. However, our previous studies demonstrated that HIV-1 Gag enters the nucleus and binds to unspliced HIV-1 RNA at transcription sites, suggesting that genomic RNA selection may occur in the nucleus. In the present study, we observed nuclear entry of HIV-1 Gag and co-localization with unspliced viral RNA within 8 hours post-expression. In CD4+ T cells (J-Lat 10.6) treated with latency reversal agents, as well as a HeLa cell line stably expressing an inducible Rev-dependent provirus, we found that HIV-1 Gag preferentially localized with histone marks associated with enhancer and promoter regions of transcriptionally active euchromatin near the nuclear periphery, which favors HIV-1 proviral integration sites. These observations support the hypothesis that HIV-1 Gag hijacks euchromatin-associated histones to localize to active transcription sites, promoting capture of newly synthesized genomic RNA for packaging.

## Introduction

Retroviruses reverse transcribe their RNA genomes (gRNA) into double-stranded proviral DNA that stably integrates into the host chromosome. Following integration, which typically occurs near the periphery of the nucleus^1–3^, host transcription machinery is used to generate nascent unspliced viral RNA (USvRNA) that is selected by Gag and packaged into new virions (as reviewed^4,5^). The retroviral Gag polyprotein is the major structural polyprotein and contains several conserved domains, including matrix (MA), capsid (CA), nucleocapsid (NC), and p6, which are individually cleaved during virus maturation (as reviewed^6^). To select gRNA for incorporation into assembling virions, the NC domain of Gag recognizes and selectively binds to the psi packaging signal positioned at the 5’ end of the gRNA transcript, forming a ribonucleoprotein complex (as reviewed^5,7^). Loss of the NC domain diminishes Gag’s ability to package gRNA into new virus particles and decreases virus-like particle (VLP) production^8^. Although the binding of Gag to gRNA has been well-characterized using *in vitro* and cell-based studies, the mechanisms that govern the gRNA selection process have been reexamined with the discovery of Gag nuclear trafficking ^9–11^. Gag proteins of various retroviruses, including human immunodeficiency virus type-1 (HIV-1)^9–11^, Rous sarcoma virus (RSV)^10,12–17^, and mouse leukemia virus (MLV)^18^, have all been found to undergo nuclear localization. For RSV, disruption of Gag nuclear trafficking decreases packaging of gRNA into virions^15^, indicating that efficient genome packaging depends not only on NC binding but also on nuclear trafficking of Gag. In both RSV and HIV infection, Gag associates with newly synthesized USvRNA at proviral integration sites, suggesting that Gag traffics to sites of active transcription^11,14^. However, it remains unclear whether Gag localizes to specific structures in the nucleus, and the timing of HIV-1 viral ribonucleoprotein complex (vRNP) formation is undefined. In this work, we examined the temporospatial properties of HIV-1 Gag localization to subnuclear structures and its association with USvRNA in the nucleus.

The nucleus is the control center of the cell where genome replication, transcription, and splicing occur. These processes are highly compartmentalized into specific subnuclear domains, including chromosome territories and nuclear splicing speckles, enabling the cell to efficiently coordinate the expression of thousands of genes (as reviewed^19^). Aside from subnuclear compartments, chromatin organization and genome architecture also play a major role in gene regulation and cellular differentiation (as reviewed^20,21^). In HIV-1, the reverse-transcribed proviral DNA enters the nucleus to be loaded with histones and then preferentially integrates into active gene locations near the nuclear periphery^1–3,22^. The transition between active and latent HIV-1 infection is regulated by the chromatin state^23–25^. Condensation of the chromatin surrounding the HIV-1 integrated provirus promotes transcriptional repression, allowing the establishment of latency and creating a long-lived reservoir of infected T cells (as reviewed^26^). This series of events permits the virus to remain undetected and evade host immune responses. Accordingly, recent efforts to combat HIV-1 infection, such as the “shock and kill” method, aim to reverse HIV-1 latency through the use of latency reversal agents (LRAs) to trigger cell death in infected T cells^27–30^.

One factor that influences HIV-1 transcriptional control is epigenetic modifications that alter the chromatin structure. For example, H3 and H4 acetylation are associated with increased HIV-1 transcription through the recruitment of histone acetyltransferases to the proviral long terminal repeats (LTR)^31–33^. Conversely, H3 methylation leads to HIV-1 transcriptional repression by restricting LTR access and promoting heterochromatin formation with sequestration of HIV-1 Tat^34–40^. Together, these studies demonstrate that the spatial properties of nuclear proteins and chromatin architecture can influence HIV-1 gene expression.

Learning more about the spatial biology and temporal distributions of the nuclear population of HIV-1 Gag is important to uncover novel functions of Gag that have yet to be explored. Many fundamental questions about the mechanism and role of HIV-1 Gag nuclear trafficking remain unanswered. We recently showed that co-localization of HIV-1 Gag and USvRNA increases in cells treated with the transcription inhibitors actinomycin D and DRB, suggesting that Gag binds USvRNA co-transcriptionally^11^. As a follow up to those studies, in this work we examined whether HIV-1 Gag traffics to areas of the nucleus enriched in euchromatin, based on its localization to sites of active vRNA transcription.

## Materials and Methods

### Plasmid, Cells, and Inductions

HIV-1 Gag-CFP rtTA HeLa cells contain a doxycycline-inducible proviral plasmid stably integrated into a HeLa cell line containing rtTA as previously described^11^. Briefly, the original proviral plasmid was constructed from the HIV-1 NL4.3 strain containing 24 copies of the MS2-MBL stem loops^41^ (a kind gift from Dr. Nathan Sherer, University of Wisconsin-Madison). The plasmid was modified by deletion of 21 MS2-MBL loops, substitution of the HIV-1 LTR with a dox-inducible promoter at both the 5’ and 3’ ends of the provirus, excision of a 1.9 kB region of the *pol* gene using *MscI* sites, and replacement of the *nef* gene with rtTA. The resulting DNA was cloned into the Piggybac vector (System Biosciences, Palo Alto, CA, USA) containing a selective marker for puromycin drug resistance. A stable cell line was created using selection with puromycin (Cytiva HyClone, Malborough, MA, USA). The HeLa cell line was maintained in Dulbecco’s modified Eagle medium containing 10% fetal bovine serum. Expression of the provirus was induced by treating cells with doxycycline (GoldBio, St. Louis, MO, USA) at various concentrations, as stated in the Results section. J-Lat T cells (clone 10.6, obtained through the NIH AIDS Reagent Program) were maintained in RPMI medium supplemented with 10% fetal bovine serum. Both cell lines were grown in the presence of penicillin, streptomycin, and fungizone at 38°C and 5% CO_2_. Reversal of latent HIV-1 gene expression in J-Lat 10.6 cells was performed by treatment with prostratin (Sigma, St. Louis, MO, USA) or TNFα (Abcam, Cambridge, MA, USA), as noted in the Results section.

### Subcellular Fractionation

Fractionation of HIV-1 Gag-CFP rtTA HeLa cells was performed as described with minor modifications^11^. Briefly, cells were grown in 10 cm^2^ cell culture dishes (Genesee Scientific GenClone, San Diego, CA, USA), doxycycline induced, trypsinized, washed with cold 1x phosphate saline buffer (PBS), and centrifuged at 16,000 x *g* at 4°C for 30 sec to remove residual culture medium. Cells were resuspended in 400 µL cold cytoplasmic lysis buffer (10 mM HEPES pH 7.9, 10 mM KCl, 0.1 mM EDTA, 0.4% [vol/vol] Nonidet P40, 1 mM DTT, Roche protease inhibitor cOmplete tablets EDTA free (MilliporeSigma, St. Louis, MO, USA) to 1×)) and incubated on ice for 5 min, followed by centrifugation at 16,000 x *g* at 4°C for 5 min. Supernatants were collected as the cytoplasmic fraction and pellets were resuspended in 200 µL cold nuclear lysis buffer (20 mM HEPES pH 7.9, 100 mM NaCl, 1 mM DTT, Roche protease inhibitor cOmplete tablets EDTA free) in the presence of OmniCleave (Biosearch Technologies, Novato, CA, USA). Pellets were incubated in a 37°C water bath for 10 min and cell suspensions were brought up to 400 mM NaCl and 1 mM EDTA. Pellets were vortexed for 15 sec and rotated end-over-end at 4°C for 20 min. Samples were spun at 16,000 x *g* at 4°C for 10 min and the supernatant were collected as the nuclear fraction.

Fractionation of J-Lat 10.6 cells were performed similarly as described above with minor modifications for the cytoplasmic lysis. Prior to lysis, J-Lat 10.6 cells were grown in T75 flasks (Corning Inc., Corning, NY, USA) and induced by prostratin or TNFα treatment. Cells were resuspended in 1000 µL cold cytoplasmic lysis buffer (10 mM HEPES pH 7.9, 10 mM KCl, 2 mM Mg acetate, 3 mM CaCl_2_, 340 mM sucrose, 1 mM DTT, Roche protease inhibitor cOmplete tablets EDTA to 1×) and incubated on ice for 5 min. Nonidet P40 (0.05%) was added to samples and immediately vortexed for 15 sec followed by centrifugation at 3,500 x *g* at 4°C for 10 min. Supernatants were collected as the cytoplasmic fraction. Nuclear fractions were extracted as described above.

Chromatin fractionation of HIV-1 Gag-CFP rtTA HeLa and J-Lat 10.6 cells were performed as described by Chase et al^42^. Following induction and harvest, cells were resuspended in 1000 µL cold sucrose buffer containing (10 mM HEPES pH 7.9, 10 mM KCl, 2 mM Mg acetate, 3 mM CaCl_2_, 340 mM sucrose, 1 mM DTT, Roche protease inhibitor cOmplete tablets EDTA to 1×) and incubated on ice for 5 min, followed by centrifugation at 3,500 x *g* at 4°C for 10 min. The supernatant was collected as the cytoplasmic fraction. Cell pellets were then resuspended in 1000 µL cold nucleoplasm extraction buffer (50 mM HEPES pH 7.9, 150 mM K Acetate, 1.5 mM MgCl_2_, 0.1% [vol/vol] Nonidet P40, 1 mM DTT, Roche protease inhibitor cOmplete tablets EDTA free) and homogenized using Dounce homogenizers on ice. Samples were transferred into fresh tubes and rotated end-over-end at 4°C for 20 min. Supernatant was collected as the nucleoplasm fraction after centrifuging samples at 16,000 x *g* at 4°C for 10 min. Remaining cell pellets were resuspended in 500 µL cold nuclease incubation buffer (50 mM HEPES pH 7.9, 10 mM NaCl, 1.5 mM MgCl_2_, 1 mM DTT, 100 U/mL Omnicleave (Biosearch Technologies, Novato, CA, USA), Roche protease inhibitor cOmplete tablets EDTA free) and incubated in a 37°C water bath for 10 min. The buffer in cell suspension was adjusted to 150 mM NaCl and incubated on ice for 20 min. Samples were centrifuged at 16,000 x *g* at 4°C for 10 min and the supernatant was collected as the euchromatin fraction. Cell pellets were then resuspended in 500 µL cold chromatin extraction buffer (50 mM HEPES 7.9, 500 mM NaCl, 1.5 mM MgCl_2_, 0.1% Triton X-100, 1mM DTT, Roche protease inhibitor cOmplete tablets EDTA free) and incubated on ice for 20 min. Samples were centrifuged at 16,000 x *g* at 4°C for 10 min and the supernatant was collected as the heterochromatin fraction. Chromatin fractionations in experiments involving the treatment with HDACi romidepsin (Sigma, St. Louis, MO, USA) were performed as described above with 18 nM romidepsin treatment prior to fractionations.

### Western Blot Analysis

After cell fractionation, protein concentrations were determined by Bradford Protein Assay protein using Coomassie protein assay reagent (Thermo Fisher Scientific-Invitrogen, Waltham, MA, USA). 5 µg of each protein fraction was added to 1x loading buffer (250 mM Tris pH 6.8, 40% glycerol, 8% β-mercapto-ethanol, 0.4% bromophenol blue, 8% SDS) and subjected to immunoblot analysis using the Stain-Free Imaging Technology (BioRad, Hercules, CA, USA). After SDS-Page, gels were transferred to PVDF membrane and blocked in 5% milk TBST (1×Tris-buffered saline, 0.1% Tween-20) for 30 min. Gag was detected using purified mouse anti-p24 supernatant from anti-HIV-1 Gag hybridoma 183 (NIH AIDS Reagent Program, Division of AIDS, NIAID, NIH, Germantown, MD, USA) at 1:2000 dilution. Antibodies targeting Calnexin (1:3000, Enzo, Cat #: ADI-SPA-865-F), Med4 (1:5000, Abcam Cat #: Ab129170), H3K4me3 (1:5000, Abcam, Cat #: Ab8580), and H3K27ac (1:2500, Abcam, Cat #: Ab4729) were used to assess fraction purities. Mouse (1:5000) and Rabbit (1:10,000) secondary HRP-conjugated antibodies (Jackson ImmunoResearch, West Grove, PA, USA; mouse Cat #: 711-036-152; rabbit Cat #: 715-036-150) were used with Pierce ECL Western Blotting Substrate (Thermo Scientific, Cat #: 32106) and visualized by a ChemiDox MP Imager (BioRad). Densitometry analysis was performed using ImageLab (BioRad) looking at lane statistics. Adjusted total band volume data was extracted for analysis.

Meta-analysis of Gag localization in cytoplasmic and nuclear fractions was performed by normalizing to total protein of each protein fraction. Briefly, total protein amounts were determined using the protein concentration acquired by the Bradford Protein Assay. Next, we calculated the percent 5 µg of protein represented and determined a normalization factor to equal 1% of total protein for each fraction to adjust for differences in the total protein of the cytoplasmic and nuclear fractions. Adjusted total band volume, which represents the 5 µg of protein loaded on the western blot, was adjusted by the normalization factor to calculate the band volume representing 1% of the total protein. Newly adjusted cytoplasmic and nuclear band volumes were used to calculate the percent Gag localization. Data quantitating the percent of nuclear Gag and the percent of chromatin marks in nuclear fractions were statistically analyzed through two-way ANOVA and one-way ANOVA, respectively.

### Single molecule Fluorescence in-situ Hybridization (smFISH)

HIV-1 Gag-CFP rtTA HeLa cells were induced with doxycycline (2 µg/mL) for 24 hrs. Cells were washed with PBS and fixed with 3.7% formaldehyde in 1x PBS for 10 min. Single molecule FISH (smFISH) was performed using Stellaris™ RNA FISH protocol in Adherent Cells (Biosearch Technologies, Novato, CA, USA). After fixation, cells were washed twice with 1x PBS and permeablized with 70% EtOH overnight at 4°C, then incubated in rehydration buffer (10% formamide, 2x SSPE, DEPC water) for 20 min at room temperature. Coverslips with cells were inverted onto RNA FISH probes diluted 1:50 in hybridization buffer (10% dextran, 10% formamide, 2x SSPE, DEPC water) in a humid chamber overnight at 37°C. Coverslips were then washed with rehydration buffer for 30 min at 37°C followed by DAPI staining (1:1000) for 30 min. Coverslips were mounted onto microscope slides using ProLong Diamond Antifade Mountant (Thermo Fisher Scientific-Invitrogen, Waltham, MA, USA) and cured for 72 hrs.

### Immunofluorescence of Histone Markers

HIV-1 Gag-CFP rtTA HeLa cells were briefly washed with PBS and fixed using 3.7% formaldehyde for 10 min at room temperature. Following fixation, cells were washed 3 times for 5 min with PBS and permeabilized with permeabilization buffer (0.12 M KCl, 0.02 M NaCl, 0.01M Tris-HCl pH 7.7, 0.1% Triton X-100) for 10 min. Cells were washed again three times and blocked with 3% BSA in PBS for 10 min. Primary antibodies against H3K4me3 (1:600; Abcam, Cat #: Ab8580), H3K27ac (1:600; Abcam, Cat #: Ab4729), and H3K9me3 (1:600, Active Motif, Cat #: 39766) were diluted in blocking buffer and applied to inverted coverslip containing cells for 45 min at room temp. Cells were then washed 3 times using blocking buffer for 5 min each. AlexaFluor 647 secondary antibodies (1:1000; Life Technologies, Cat #: A31573) were applied similarly as primary antibody for 45 min at room temp. Cells were then washed three times with PBS again and mounted onto microscope slides using ProLong Diamond Antifade Mountant (Thermo Fisher Scientific-Invitrogen, Waltham, MA, USA). Following preparations of slides, cells were imaged by confocal microscopy and analyzed through Imaris 10.0 image analysis software (Bitplane Inc., Concord, MA, USA). Data recorded were statistically tested using one-way ANOVA for significance.

### Confocal Microscopy

Following preparation of cells, coverslips were imaged using a Leica SP8 laser scanning confocal microscope (Leica Microsystems, Buffalo Grove, IL, USA). Imaging was performed by a white light laser using a 63X/NA1.4 oil immersion objective lens with sequential scanning of each channel with a frame average of 4 and at 400 Hz imaging speed. Single fluorophore controls were imaged separately to ensure no cross talk occurred. Images for the HIV-1 Gag time course experiments were processed by Gaussian filters and histograms were adjusted to best visualize nuclear foci. Images of Gag and histone marker co-localization were processed by deconvolution using Huygens Image Processing Software (Scientific Volume Imaging). Three-dimensional surface renderings of Z-stacks, cross-sections, and co-localization channels were generated in Imaris 10.0 image analysis software (Bitplane).

Time-lapse movies of images captured by confocal microscopy were created using Imaris image analysis software (Bitplane). For FISH experiments presented in Fig. 2, cells were rotated along the X-plane in 360° followed a scanning through the Z-plane using the Clipping Plane function. Movies created for experiments involving the visualization of histone markers (Fig. 6) were done by first rotating the cells along the X-plane in 360° and then the Y-plane in 90°. Cells were zoomed in to visualize the co-localized Gag:histone spots. All movies were presented at 15 frames per sec speed.

Movies of images captured by confocal microscopy were created using Imaris image analysis software (Bitplane). For FISH experiments presented in Fig. 2, cells were rotated along the X-plane in 360° followed a scanning through the Z-plane using the Orthogonal Clipping Plane function. Movies created for experiments involving the visualization of histone markers (Fig. 6) were done by first rotating the cells along the X-plane in 360° and then the Y-plane in 90°. Cells were zoomed in to visualize the co-localized Gag:histone spots. All movies were presented at 15 frames per sec speed.

### Quantitative Image Analysis

To measure the intensity of Gag localization across cytoplasmic and nuclear compartments in the time course experiments, fluorescent intensity analysis was performed using ImageJ. Images were selected in Leica LAS X by isolating a single Z plane through the center of the cell nucleus. Nuclei were traced in ImageJ and the fluorescent intensity was measured. Background subtraction was achieved by sampling a small region outside the cell and averaged with all cell replicates to obtain the corrected total cell fluorescence.

To determine the percent of Gag and USvRNA co-localization, Imaris image analysis software (Bitplane) was used to calculate a Mander’s Coefficient based on the HIV-1 Gag and USvRNA signal from captured images. The nuclear signal for each channel was isolated by creating a mask of the nucleus generated from surface rendering of the DAPI stain. Co-localization analysis between Gag and USvRNA was performed using the co-localization function and used to generate a co-localization channel, and to obtain the Mander’s coefficient. Similar methods were employed to determine the co-localization of HIV-1 Gag with various histone markers. To measure the distances from the chromatin periphery, a surface rendering of the histone markers was generated and the spot function was used to generate spots from the Gag:histone co-localization channel. The distance between Gag:histone co-localized spots and the periphery of the surface was measured.

## Results

### HIV-1 Gag nuclear trafficking occurs during early phases of Gag expression and co-localizes with USvRNA

To explore the temporospatial dynamics of HIV-1 Gag nuclear localization, we constructed a HeLa cell line containing an integrated proviral genome expressing HIV-1 Gag-CFP controlled by a doxycycline promoter (Fig. 1A). We performed a time course using the proviral HIV-1 Gag-CFP rtTA HeLa cell line by treating cells with doxycycline (2 µg/mL). Subcellular fractionation was performed every 4 hours for 24 hours to measure the amount of Gag protein present in cytoplasmic and nuclear fractions. Gag expression was detected by western blot using an anti-p24 antibody targeting the CA domain of Gag, and quantification of band intensities was measured by densitometry analysis. Our results indicated that HIV-1 Gag was present in cytoplasmic and nuclear fractions as early as 4 hours post-induction (h.p.i.; Fig. 1B and C) and expression in each fraction increased through 20 hrs, plateauing at approximately 24 hrs. Fraction purity was confirmed using antibodies against Calnexin (cytoplasm) and Med4 (nucleus). At each time point, Gag levels were statistically significantly higher in the cytoplasmic fractions compared to the nuclear fractions (Fig. 1D), with a range of approximately 30-37% of Gag in the nuclear fraction. The percent of Gag in the nucleus vs. cytoplasm remained relatively constant throughout the time course (Fig. 1D). The presence of full-length Gag in the nucleus at early time points (4-8 hours) indicates that Gag nuclear trafficking occurs shortly after Gag production begins and is not driven by a concentration gradient. Moreover, Gag levels fluctuate between the cytoplasmic and nuclear compartments over time, indicating that the nuclear population of Gag is either stable or replenished after it is depleted from the nucleus.

**Figure 1.**
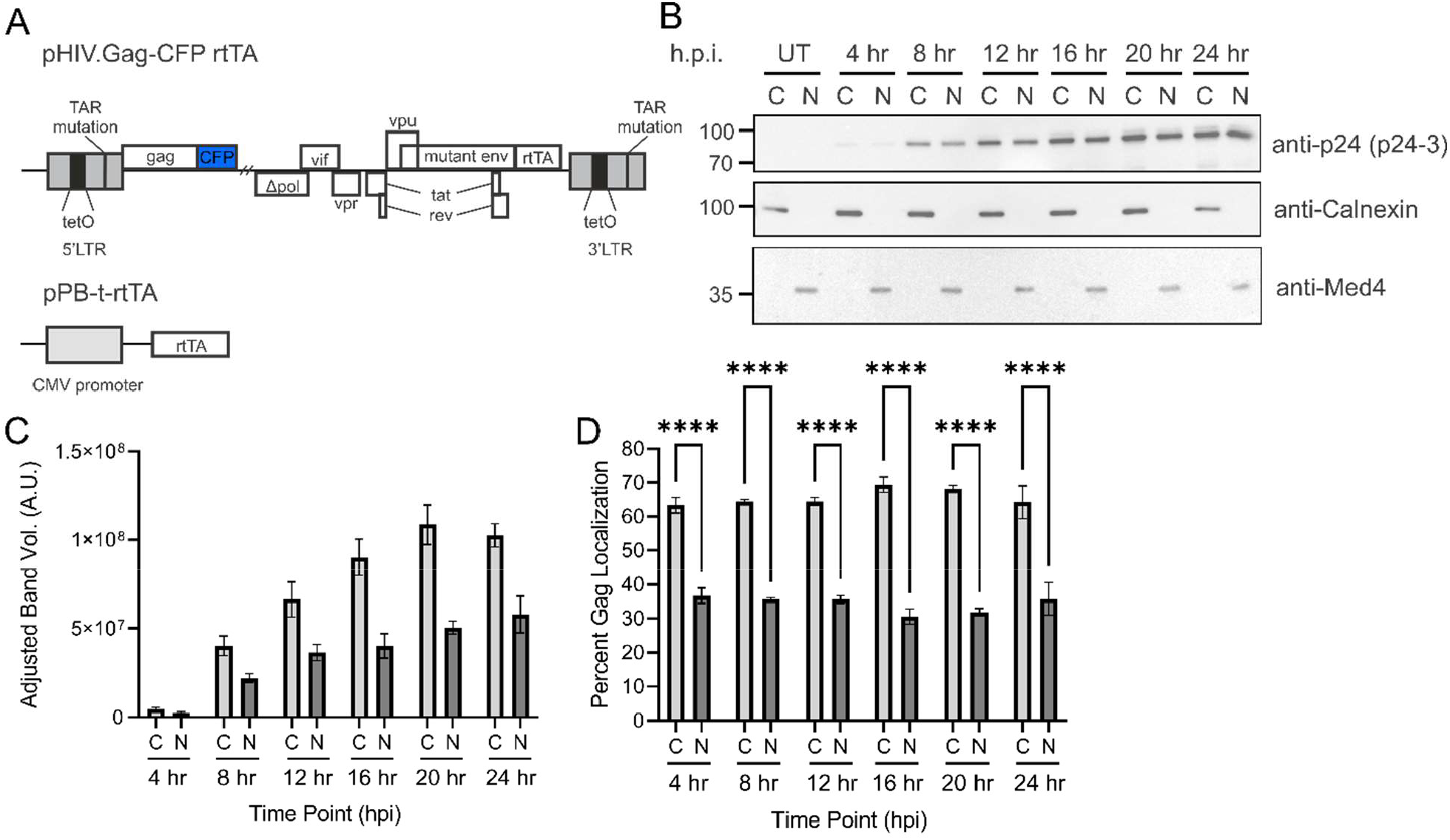
Nuclear Gag localization occurs within 4 hrs in stably-integrated HeLa cells. (A) Schematic of the doxycycline-inducible HIV-1 proviral construct expressing Gag-CFP and rtTA plasmids integrated in HeLa cells (HIV-1 Gag-CFP rtTA HeLa). The proviral construct contains a deletion in *pol*, frameshift mutation in *env*, and substitution of *nef* with rtTA. The 5’ LTR contains a TAR mutation and insertion of TetO sites at the promoter. (B) HIV-1 Gag-CFP rtTA HeLa cells were induced with dox (2 µg/mL) in a time-dependent fashion every 4 hours and subcellular fractionated to isolate cytoplasmic and nuclear lysates. Western blotting was performed to observe full-length HIV-1 Gag. Fraction purity was assessed by Calnexin (cytoplasmic) and Med4 (nuclear). (C) Densitometry of immunoblots of cytoplasmic and nuclear fractions at each time point shown as adjusted band volumes. Background subtraction was performed to obtain adjusted band volume. (D) Percent Gag localization was determined by (fraction band volume)/(total band volume of cytoplasmic and nuclear within each time point). Analysis was compiled over 3 independent replicates (error bars = standard error of mean; statistical significances: **** *P* < 0.0001).

To examine HIV-1 Gag-CFP subcellular localization using an imaging approach, several time points were imaged using confocal microscopy (8, 12, 16, 20, and 24 h.p.i.). Beginning at 8 h.p.i, both focal and diffuse Gag-CFP fluorescence (green) was observed within the nucleus (nuclear DNA stained blue by DAPI; Fig. 2A), in the cytoplasm, and at the plasma membrane. At each time point, Gag and USvRNA co-localized in the nucleus (yellow; Merged image panels in Fig. 2A). Co-localization was also observed for distinct Gag:USvRNA complexes in the cytoplasm and at the plasma membrane. Three-dimensional reconstruction of Z-stacks confirmed Gag:USvRNA co-localization in the nucleus in each plane as indicated by the arrow (white; co-localization channel). Additionally, a surface rendering of the nucleus and co-localized spots were created to confirm co-localization was occurring within the nucleus (8 and 24 h.p.i.; Fig. 2B). Analysis of three-dimensional renderings of individual Z-stacks showed Gag and USvRNA co-localization within the nucleus for the 8 and 24 h.p.i. time points (Movie 1 and Movie 2, respectively).

To analyze nuclear Gag levels obtained by confocal imaging, nuclear Gag fluorescence was measured from each single Z-plane (n=30-41 cells). Comparison of Gag levels in the nuclear compartments yielded similar trends as observed in Fig. 1, with more Gag in the cytoplasm compared to the nucleus. The percent of nuclear Gag decreased from a peak at 8 h.p.i. (Fig. 2C). It should be noted that there was a lower ratio of nuclear:cytoplasmic ratio of Gag detected using fluorescence imaging compared to western blotting. This discrepancy may be attributed to several factors. One factor is that the population size these analyses were different; the fractionation and western blot analysis represented a large population of cells (n≈9 million cells) whereas imaging was performed on a single-cell basis and represented a smaller population (n=30-41 cells). Because the HIV-1 Gag-CFP rtTA HeLa is a polyclonal cell line and although all cells were treated with doxycycline, proviral transcription levels and Gag expression varied from cell to cell. Another factor is the method and sensitivity of protein detection, comparing chemiluminescence of western blots versus CFP epifluorescence signal in confocal imaging. The higher sensitivity of Gag detection in western blot could be explained by the ability to detect monomeric Gag more efficiently than in imaging of CFP, which does not detect single molecules or faint Gag foci. Furthermore, it is important to note that the imaging analysis was performed by analyzing a single Z slice through the center of the nucleus, which does not account for Gag molecules in the three-dimensional volume of the nucleus. Bearing in mind the discrepancies in absolute amounts, the overall trend of nuclear Gag localization is similar between both methods.

In addition, single cell images taken for each time point containing HIV-1 Gag and USvRNA were subjected to quantitative co-localization analysis using Imaris (n=9-26 cells) by measuring the overall fluorescence intensity for Gag and USvRNA signals. At 8 h.p.i., we observed a lower percentage of nuclear Gag co-localized with USvRNA (11.9%; Fig. 2D) and nuclear USvRNA co-localized with HIV-1 Gag (13.5%; Fig. 2E). By 12 h.p.i., the percentage of nuclear USvRNA that was co-localized with Gag increased (26.5%), followed by a decrease in co-localization through the 24-hour time point (22.6%; Fig. 2E). The percentage of Gag co-localized with USvRNA increased after 8 h.p.i. and remained relatively constant by 16 h.p.i. (25.5%; Fig. 2D). The overall percentage of HIV-1 Gag co-localized with USvRNA was similar to that previously reported by Tuffy et al^11^, although a different analytical method (object-based co-localization) was used in that study.

### HIV-1 Gag preferentially localized to the euchromatin-associated protein fraction

Our previous results suggested that Gag localizes to the proviral transcription site in HeLa and T cells^8^. Therefore, we examined whether Gag trafficked to specific chromatin-associated regions. Initially, we tested whether the level of HIV-1 Gag protein expression affected its localization properties. HIV-1 Gag-CFP rtTA HeLa cells were treated with varying concentrations of doxycycline to regulate the level of Gag expression. After isolation of nuclei, extraction of euchromatin- and heterochromatin-associated proteins was achieved using different salt concentrations (150 nM NaCl and 500 nM NaCl, respectively) and western blotting was performed to measure the level of Gag present in the euchromatin (Ch150), and heterochromatin (Ch500) protein fractions (Fig. 3A). Purity of each fraction was assessed using antibodies against Calnexin (cytoplasm), Med4 (nuclear), and tri-methylated H3K4 histone (H3K4me3, euchromatin). H3K4me3 has previously been shown to mark active promoters across different cell types^43,44^. As reported by Chase et al.^42^, Ch150 extracted protein fractions contained active cellular transcription proteins that were distinct from the Ch500 fraction despite the presence of H3K4me3 in both chromatin fractions^42^. Densitometry analysis of western blots revealed that HIV-1 Gag preferentially localized with euchromatin (69.1%) compared to heterochromatin (30.9%) at lower Gag expression levels (20 ng/mL doxycycline) (Fig. 3B). Similar results were seen using 200 ng/ml doxycycline induction, with 58.1% of Gag in the euchromatin and 41.9% in heterochromatin fraction. At the highest doxycycline induction level, the ratio of Gag in euchromatin vs. heterochromatin fractions was approximately 50:50. To ensure the treatment of doxycycline did not alter the chromatin, H3K4me3 expression was measured and found to be similar across each drug concentration treatment (Fig. 3C).

**Figure 2.**
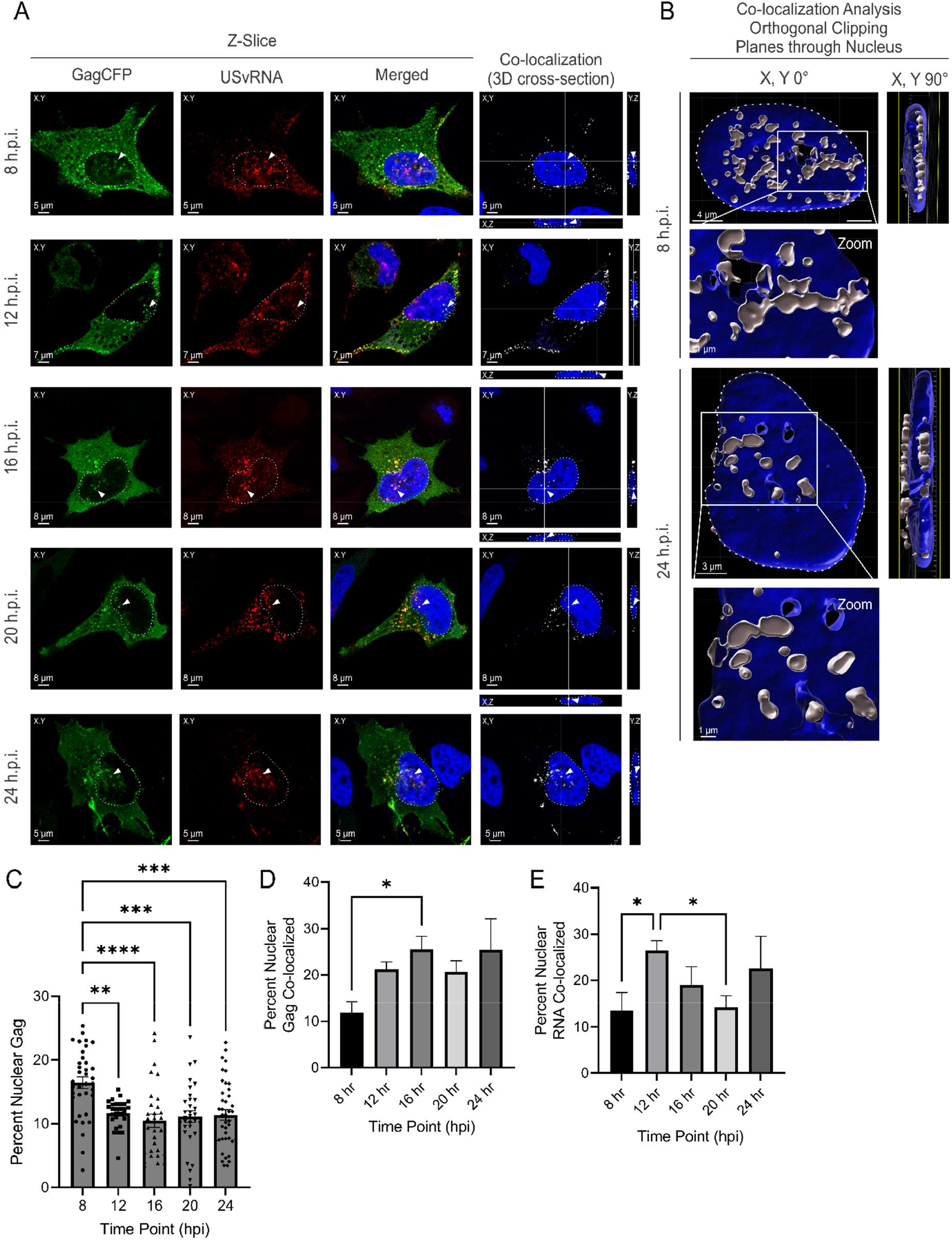
HIV-1 Gag co-localizes with unspliced vRNA (USvRNA) shortly after Gag expression in integrated HeLa cells. (A) Confocal images of HIV-1 Gag-CFP rtTA HeLa cells at 8, 12, 16, 20, and 24 h.p.i. visualizing Gag-CFP (green) and USvRNA (red) in the XY plane. Nuclei of cells were visualized by DAPI stain (blue) and co-localization analysis identified Gag and vRNA co-localized signal (white) across XY, YX, and YZ planes. (B) Three-dimensional surface rendering depicts 8 and 24 h.p.i. time points to demonstrate that the Gag and USvRNA co-localization occurs within a single orthogonal clipping plane through the XY plane of the nucleus. Images were generated by Imaris of the center of the nucleus and rotated (90°) planes. (C) The percentage of nuclear Gag was calculated from corrected cell fluorescence of confocal images. Quantitative analysis of HIV-1 Gag-CFP rtTA HeLa cells induced with doxycycline measuring nuclear HIV-1 Gag co-localization with USvRNA (D) and USvRNA co-localization with Gag (E) by Mander’s coefficient (error bars = standard error of mean; statistical significance: * *P* < 0.05, ** *P* < 0.01, *** *P* < 0.001, **** *P* < 0.0001).

**Figure 3.**
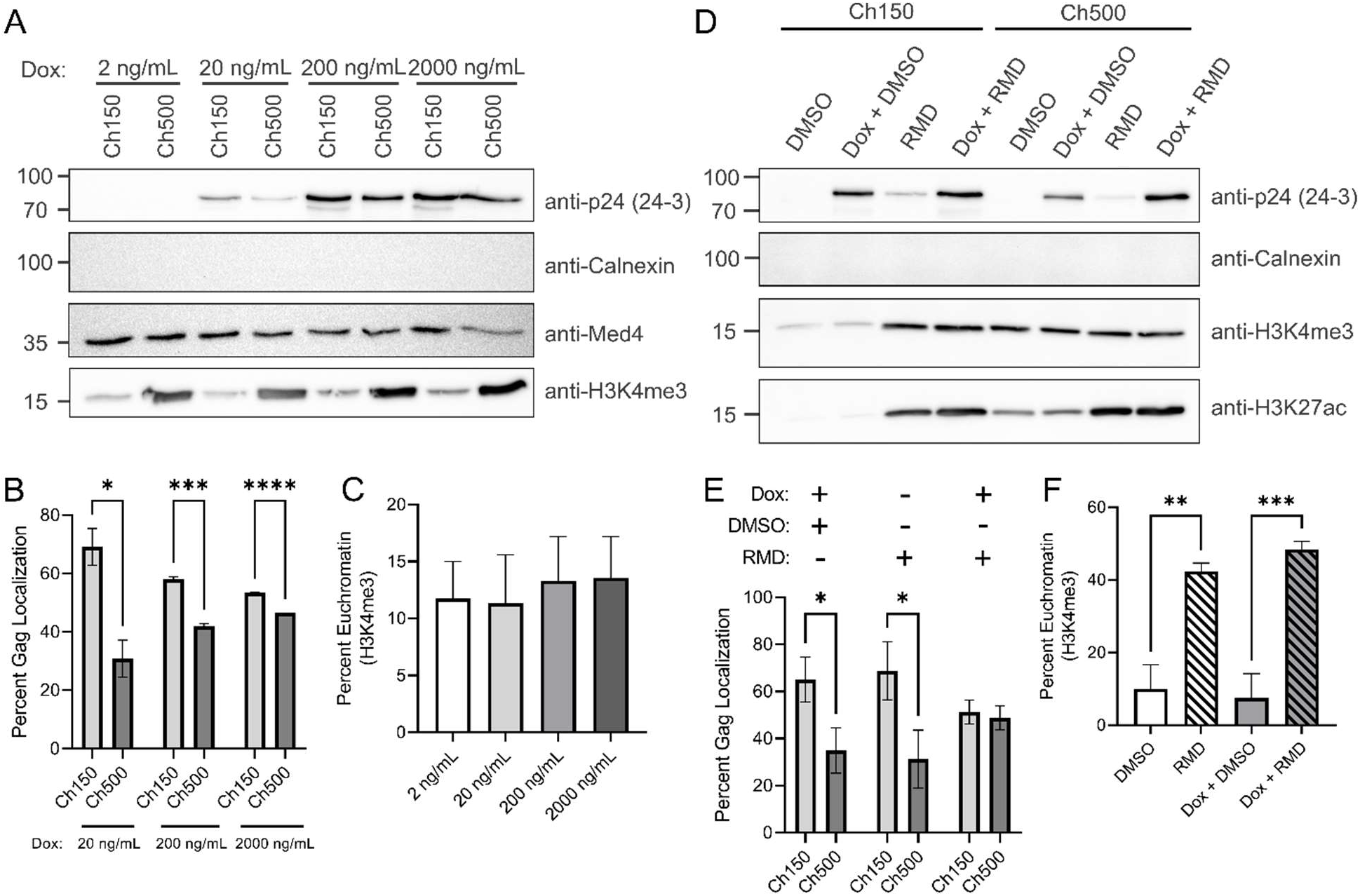
HIV-1 Gag preferentially localizes to the euchromatin protein fractions in integrated HeLa cells. (A) HIV-1 Gag-CFP rtTA HeLa cells were induced using 2 ng/mL, 20 ng/mL, 200 ng/mL and 2000 ng/mL doxycycline. Gag was detected in euchromatin (Ch150) and heterochromatin (Ch500) fractions by immunoblot using anti-p24 antibody. Calnexin (cytoplasmic), Med4 (nuclear) and H3K4me3 (chromatin) were used to assess fraction purity. (B) Analysis of detected bands from western blot measuring Percent Gag Localization. Due to the lack of signal, 2 ng/mL dox induction was excluded from the analysis. (C) Percent euchromatin was analyzed from H3K4me3 western blot signals to show no changes to euchromatin occurred during dox treatment. (D) HIV-1 Gag-CFP rtTA HeLa cell fractions following 20 ng/mL dox induction either in the presence of romidepsin (RMD) were analyzed by western blot. Anti-p24 antibodies were used to detect Gag presence while anti-Calnexin, H3K4me3, H3K27ac were used to access fraction purities. (E) The Percent of Gag Localization was determined via analysis of Gag band volumes. (F) Percent euchromatin was analyzed by H3K4me3 band volume, demonstrating that the addition of RMD increases euchromatin formation (error bars = standard error of mean; statistical significance: * *P* < 0.05, ** *P* < 0.01, *** *P* < 0.001, **** *P* < 0.0001).

We next asked whether opening the chromatin structure to be more transcriptionally-active would affect HIV-1 Gag localization. To promote euchromatin formation, cells were treated with the histone deacetylase inhibitor (HDACi) romidepsin (RMD) in the presence or absence of a low doxycycline induction (20 ng/mL) to induce proviral expression for 24 hrs (Fig. 3D). Following chromatin fractionation, analysis by western blotting determined that Gag levels in the euchromatin fraction of cells increased upon treatment with RMD (18 nM, Fig. 3E). Treatment with dox alone induced cells to exhibit higher Gag levels in euchromatin (65.1%), similar to that observed in Fig. 3B. Of note, cells treated with RMD alone (with no doxycycline addition) had higher ratios of Gag in the euchromatin (68.8%) compared to the heterochromatin fractions (31.2%; Fig. 3E). This result suggests that changing the chromatin environment around the proviral DNA increases viral gene expression independently of the doxycycline-sensitive promoter. Cells induced with doxycycline and treated with RMD displayed increased Gag expression, but with equal amounts of Gag in both chromatin fractions (Fig. 3D and E). To demonstrate the effectiveness of the HDACi drug treatment, chromatin fractions were assessed by western blotting using antibodies to H3K4me3 and acetylated H3K27 (H3K27ac) histone markers. Specifically, while H3K4me3 marks active promoters, H3K27ac marks are associated with active enhancer regions^45^. Samples treated with RMD exhibited increased H3K4me3 histone marks, signifying a shift toward euchromatin formation, as expected (Fig. 3F). Calnexin antibody was used to validate that chromatin fractions did not contain cytoplasmic contamination. These results indicate that Gag chromatin localization was influenced by the levels of Gag expressed in the cell, suggesting that the factor that allows Gag to accumulate in euchromatin may be limiting.

### HIV-1 Gag nuclear trafficking in infected T cells reactivated from latency

Based on the results observed in HeLa cells, we examined the timing and subcellular distribution of Gag in infected T cells reactivated from latency using a model system more representative of natural infection. We previously showed that HIV-1 Gag traffics to the nucleus in J-Lat 10.6 cells treated with the latency reversal agent prostratin^11^. Prostratin is a protein kinase C (PKC) agonist that acts by stimulating PKC and subsequently activating the NK-κB signaling pathway^46^. To determine the timing of Gag expression and nuclear trafficking, we performed a 24-hour time course similar to that shown in Figure 1 using J-Lat 10.6 cells. Cells were treated with prostratin (1 µg/mL) and examined every 4 hours by subcellular fractionation and western blotting. Similarly, Gag was detected in both cytoplasmic and nuclear fractions in J-Lat 10.6 cells starting at 4 h.p.i (Fig. 4A and B) with a slightly higher amount of Gag in the nucleus (58.5%) compared to the cytoplasm (41.5%; Fig. 4C). All other time points exhibited slightly higher cytoplasmic Gag, although not to a statistically significant degree. Moreover, we noticed that nuclear Gag concentrations decreased over the 24-hr period, similar to the HeLa cell model with the exception of the last time point. These subtle differences in the timing of Gag localization between the HeLa and J-Lat 10.6 cell line may be attributed to the cell type or the presence of Gag-Pol, which is absent in the HIV-1 Gag-CFP rtTA HeLa cells. Interestingly, Gag-Pol was noted to be present in the nucleus beginning at 12 h.p.i., which has not been reported previously (Fig. 4A) and warrants additional study.

**Figure 4.**
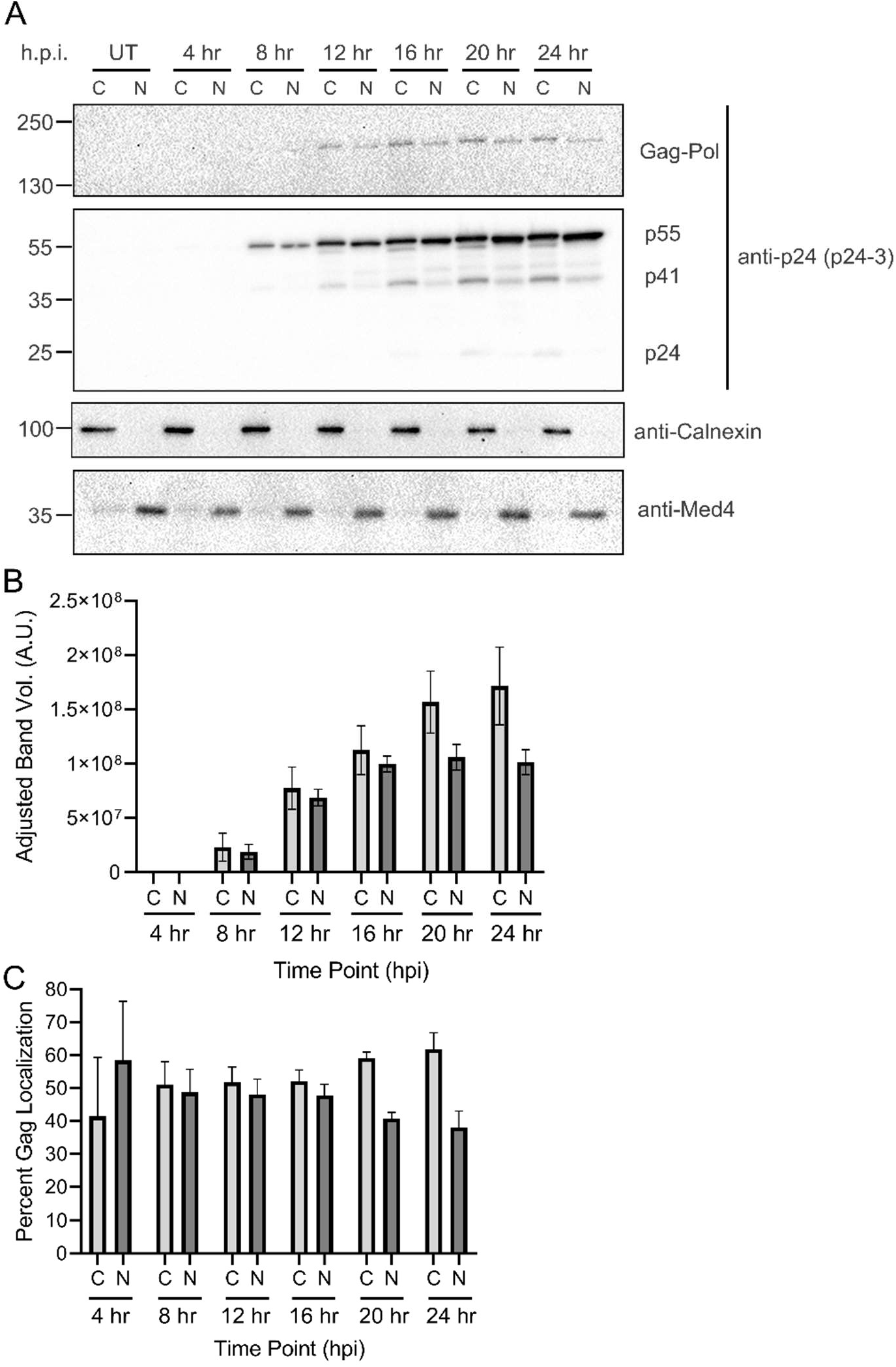
Nuclear Gag localization occurs within 8 hrs in latently-infected T cells. (A) JLAT 10.6 cells were induced with prostratin (2 µg/mL) in a time-dependent fashion every 4 hours. Cells were subjected subcellular fractionation to isolate cytoplasmic and nuclear protein fractions, and probed via western blot using anti-p24 Ab. Calnexin (cytoplasmic) and Med4 (nuclear) were used to assess fraction purity. (B) Immunoblot analysis of cytoplasmic and nuclear fractions at each time point are shown by measuring band volumes with background subtraction. (D) Percent of Gag localization was determined as Fig. 1D.

### HIV-1 Gag preferentially localized to euchromatin in latently-infected T cells treated with latency reversal agents

Previously, we reported Gag and USvRNA co-localize at proviral transcriptional burst sites in reactivated J-Lat 10.6 cells, suggesting that Gag may traffic to sites of USvRNA transcription upon reactivation of latent virus expression^11^. To determine whether Gag may be associated with the euchromatin-associated protein fraction, we assessed the chromatin localization of Gag after reactivation of HIV-1 gene expression in J-Lat 10.6 cells using varying concentrations of prostratin (Fig. 5A) and TNFα (Fig. 5D). In chronically infected T cells, HIV-1 relies on activation of the TNF receptor (TNFR) by TNFα to promote NK-κB recruitment to the HIV-1 LTR promoter^47–49^. Thus, the use of TNFα mimics natural reactivation of latently infected T cells. Densitometry analysis of western blots revealed that Gag preferentially localized to euchromatin, particularly at lower induction levels (Fig. 5B and 5E). In fractions of J-Lat 10.6 cells reactivated by 0.25 mg/mL of prostratin, we found that 57.0% of chromatin Gag localized to the euchromatin and only 43.0% localized to the heterochromatin (Fig. 5B). The ratio of Gag localization in euchromatin:heterochromatin fractions were equivalent at higher amounts of Gag expression. Similarly, J-Lat 10.6 cells reactivated with 2.5 mg/mL TNFα exhibited 58.1% of Gag localization to the euchromatin and only 41.9% to the heterochromatin (Fig. 5E). The preference of Gag localization to euchromatin over heterochromatin was observed in cells treated with 2.5, 5, or 10 mg/mL TNFα. Treatment of cells with prostratin or TNFα did not alter overall levels of euchromatin formation, as shown by the levels of H3K4me3 (Fig. 5C and 5F, respectively), indicating that the change in Gag localization was specific.

**Figure 5.**
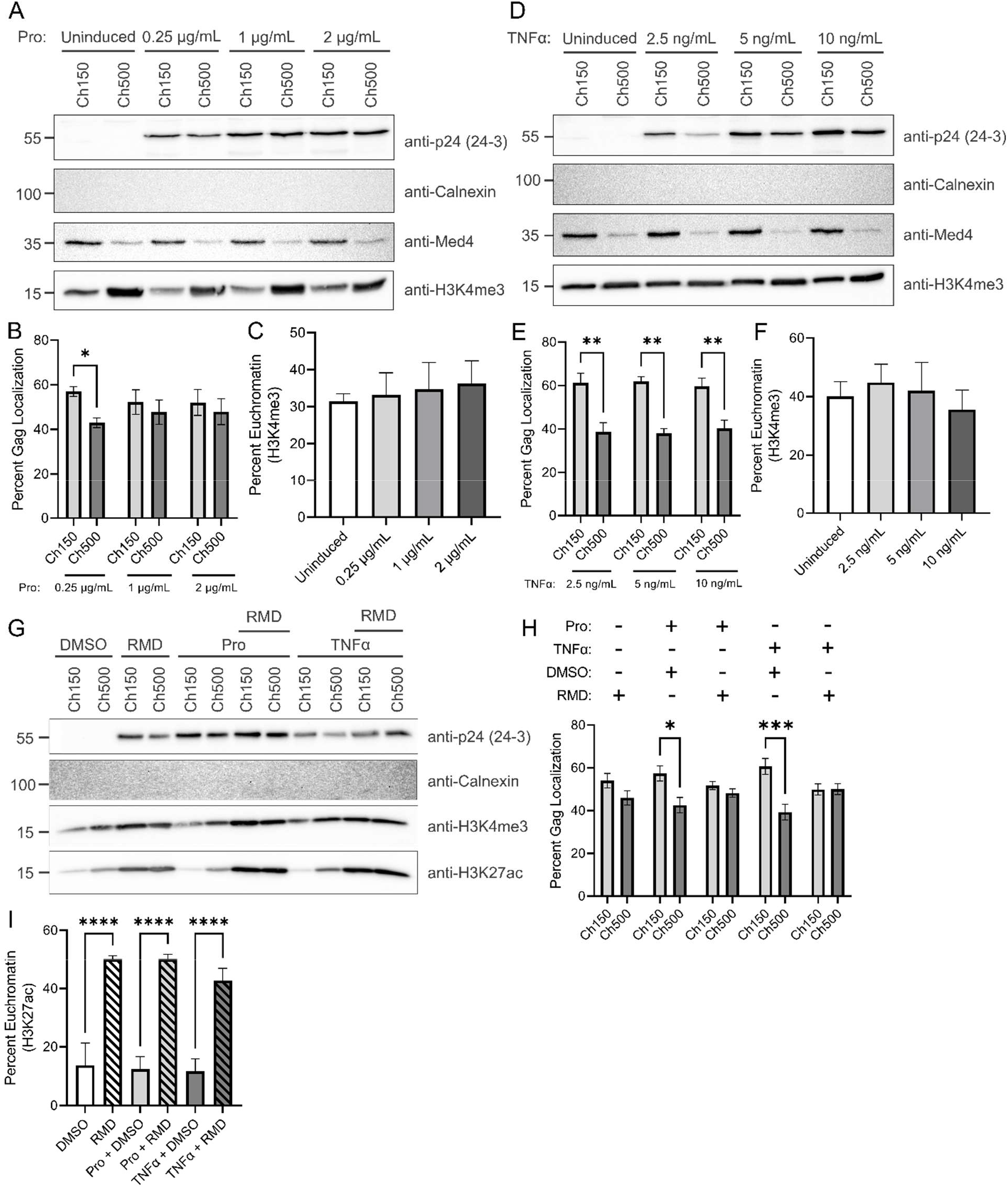
HIV-1 Gag preferentially localizes to euchromatin in latently-infected T cells. Western blot of JLAT 10.6 euchromatin (Ch150) and heterochromatin (Ch500) protein fractions following induction with various concentrations of prostratin (A) and TNFα (D). Percent Gag localization was determined from adjusted band volume from western blots of chromatin fractions following induction by prostratin (B) and TNFα (E). HIV-1 Gag is shown to preferentially localize to the euchromatin at lower inductions. Percent euchromatin was analyzed by H3K4me3 band volume for both prostratin (C) and TNFα (F) inductions. Neither induction drugs altered euchromatin formation. (G) JLAT 10.6 cells were separated into euchromatin (Ch150) and heterochromatin (Ch500) following prostratin or TNFα induction in the presence of RMD. Western blot using anti-p24 Ab was used to determine HIV-1 Gag localization. Fraction purity was assessed by antibodies targeting Calnexin (cytoplasmic), Med4 (nuclear), H3K4me3 (chromatin), and H3K27ac (chromatin). (H) Quantitation of Gag localization was determined by measuring the adjusted band volume of the western blot. (I) Assessment of percent euchromatin was analyzed through H3K27ac adjusted band volumes, demonstrating that the addition of RMD increases euchromatin formation.

The effects of HDACi on the reactivation of latently infected T cells has been extensively studied in the past, by upregulating the HIV-1 promoter and increasing Gag expression^24,30,50–52^. However, it is unknown whether HDACi would affect Gag localization and trafficking during a reactivation from latency. To determine how HDACi impacts Gag localization in T cells, we treated J-Lat 10.6 cells with RMD (10 mg/mL) coupled with either prostratin or TNFα at low induction levels (0.25 mg/mL or 2.5 mg/mL, respectively). Chromatin fractions were prepared as previously described and western blot analysis of p24 used to measure Gag levels in each nuclear fraction (Fig. 5G). Purity of each fraction was assessed using anti-Calnexin (cytoplasm), anti-Med4 (nucleus), anti-H3K4me3, and anti-H3K27ac (chromatin) antibodies. Similar to the findings shown in Fig. 5A and D, J-Lat cell induced with prostratin or TNFα alone exhibited higher Gag levels in the euchromatin (Fig. 5H). Cells treated with RMD alone also showed a preference for Gag euchromatin localization (56.1% euchromatin and 43.9% heterochromatin). However, Gag induction by prostratin or TNFα with RMD treatment resulted in a near 50:50 ratio of euchromatin and heterochromatin Gag, mimicking results seen at high levels of Gag induction. Both H3K4me3 and H3K27ac levels were increased in the presence of RMD, signifying the increased formation of euchromatin with drug treatment (Fig. 5I). Altogether, these results indicate that HIV-1 Gag preferentially localizes to the euchromatin under lower expression levels following reactivation, suggesting that localization to transcriptionally-active chromatin could be a saturable process.

### HIV-1 Gag co-localization with histone markers in the nucleus

To visualize where HIV-1 Gag localizes within the chromatin space, we performed immunofluorescence against tri-methylated H3K4 (H3K4me3), acetylated H3K27 (H3K27ac), and tri-methylated H3K9 (H3K9me3) histones (red). Following 24 hrs dox induction in our HIV-1 Gag-CFP rtTA HeLa cell line, we observed Gag co-localization with all three histone markers, independently, as indicated by the co-localization channel shown in white (Fig. 6A). Imaris co-localization analysis revealed a higher percentage of nuclear Gag co-localizes with the euchromatin markers (73.5% with H3K4me3 and 87.3% with H3K27ac) compared to the heterochromatin marker (25.9% with H3K9me3; Fig. 6B), supporting our previous observation from our chromatin fractionations (Fig. 3).

Furthermore, we assessed how far HIV-1 Gag resides from the edge of the chromatin space. To address this question, we created a surface rendering of the chromatin space (red) and used Imaris spot function to indicate where co-localization of Gag with the chromatin marks were occurring. As shown in zoomed-in panels in Fig. 6A, Gag co-localization with H3K4me3 and H3K27ac preferentially occurred near the periphery of the nucleus, extending out of the chromatin space (Movie 3 and Movie 4, respectively). Conversely, Gag and H3K9me3 co-localized foci were more buried in the three-dimensional rendering and far fewer spots were seen protruding out of the chromatin (Movie 5). Additionally, we measured the distance between Gag-histone co-localized spots and the periphery of the chromatin space, as defined by the fluorescence signal of the histone channel. Interestingly, we found that Gag co-localized with the euchromatin markers (H3K4me3 and H3K27ac) within a mean of 0.11 ± 0.002 µm of the periphery (Fig. 6C). Strikingly, co-localization of Gag with the heterochromatin histone H3K9me3 was significantly further away from the outer border of the chromatin (approximately 0.22 ± 0.002 µm, Fig. 6C), indicating that HIV-1 Gag more closely associated with the euchromatin near the periphery of the chromatin space compared to the heterochromatin.

**Figure 6.**
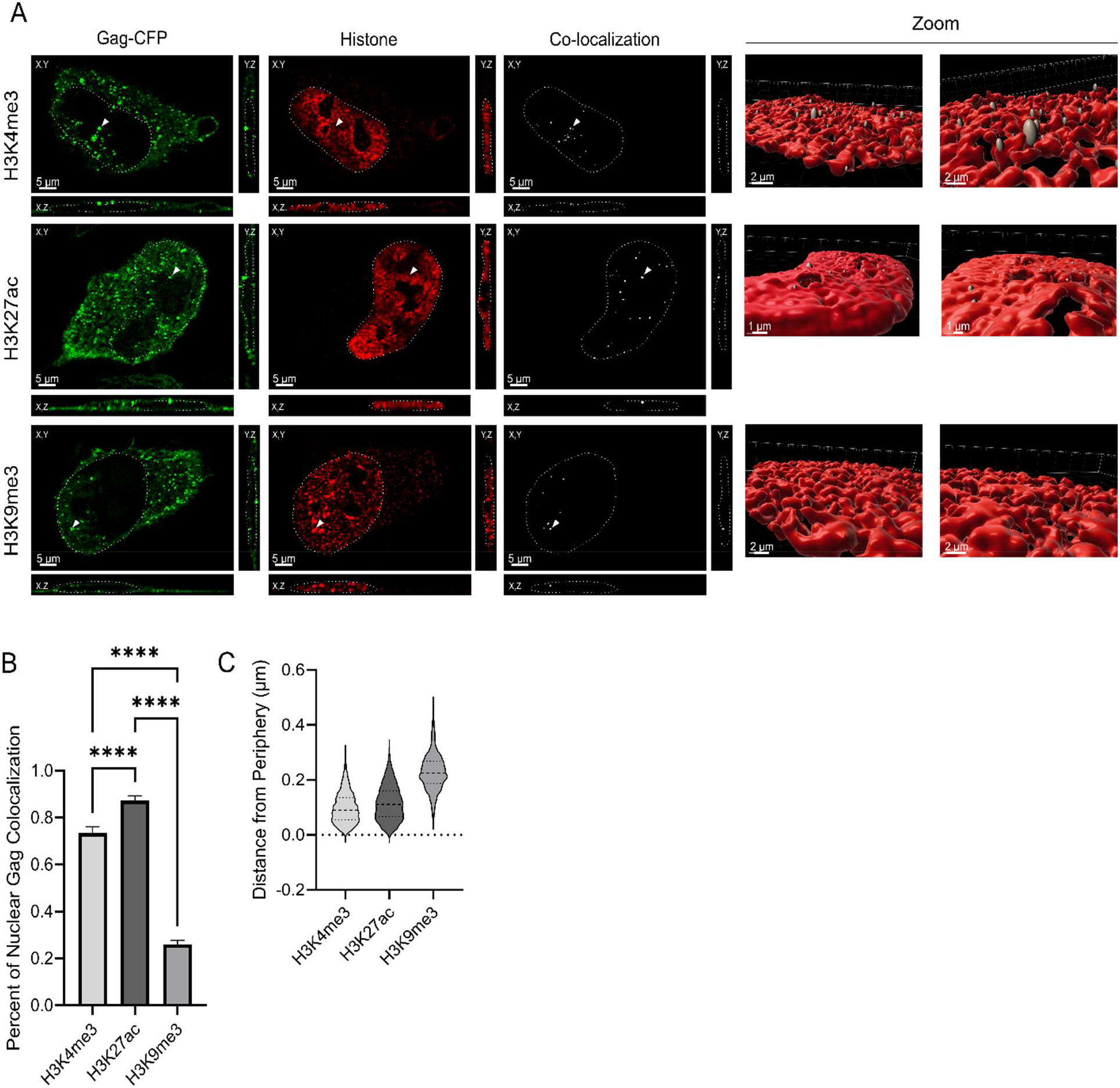
HIV-1 Gag co-localizes with euchromatin histone markers at the nuclear periphery. (A) Immunofluorescence of euchromatin (H3K4me3 and H3K27ac) and heterochromatin (H3K9me3) histone marker (red) in HIV-1 Gag-CFP rtTA HeLa cells were imaged via confocal microscopy. Images revealed that HIV-1 Gag (green) co-localizes (white channel) with each histone marker (white arrows) in the XY, XZ, and YZ planes. The chromatin space is outlined in white dashed lines. To examine the co-localized spots, a three-dimensional surface rendering of each chromatin mark was generated in Imaris and the Gag:histone co-localized signals were highlighted by the spot function. (B) The Mander’s coefficients of Gag co-localization with H3K4me3, H3K27ac, and H3K9me3 were calculated using the Imaris co-localization function. (C) The distance of Gag:histone co-localized spots to the periphery of the chromatin surface rendering was measured using Imaris (error bars = standard error of mean; statistical significance: **** *P* < 0.0001).

## Discussion

The main role of HIV-1 Gag is to bind and select USvRNA to serve as the gRNA for packaging into nascent virions. Historically, it was widely accepted that HIV-1 Gag resided only in the cytoplasm, and that the initial Gag-gRNA interaction occurred at the plasma membrane. However, with more sensitive light microscopy and advances in biochemical techniques, HIV-1 Gag has been observed in the nuclei of infected cells^10,11^. Furthermore, HIV-1 Gag was found to co-transcriptionally associate with USvRNA in the nucleus^11^, suggesting that Gag selection of USvRNA may occur earlier in the assembly pathway. Proteomic analysis of Gag interacting proteins revealed association with several nuclear factors involved in transcription and chromatin remodeling^53–57^ (Rice et al., unpublished). Together, these results suggest that HIV-1 Gag may have a novel function in the nucleus that is linked to gRNA packaging.

Using a HeLa cell line containing a dox-inducible, Rev-dependent proviral construct expressing Gag-CFP, we found that Gag expression occurred within 4 h.p.i. in both cytoplasmic and nuclear fractions (Fig. 1B and C). This property was also observed in latently infected J-Lat 10.6 cells with Gag expression beginning at 8 h.p.i. (Fig. 4A and B). A previous study by Mohammadi et al. demonstrated that USvRNA transcripts arise 8 hrs after integration and correlate with the onset of Gag protein translation^58^. Our experiments in J-Lat 10.6 cells followed a similar course, with the timing of reactivation of proviral gene expression induced by prostratin or TNFα in alignment with previous reports^58^. Given that translation of the viral mRNA Gag occurs shortly after transcription, these data suggest that Gag nuclear translocation occurs early after Gag is synthesized.

We previously showed that HIV-1 Gag co-localizes with USvRNA in the nucleus at 24 h.p.i.^11^. In the present study, we observed Gag nuclear localization as early as 4 h.p.i., therefore we sought to determine how early Gag was detected co-localizing with USvRNA. Using single-molecule RNA FISH, we observed Gag and USvRNA co-localization in the nucleus beginning at 8 h.p.i. in the HIV-1 Gag-CFP HeLa cell line (Fig. 2A). The proportion of HIV-1 Gag co-localized with USvRNA appeared to remain constant up to 16 h.p.i., whereas the percentage of USvRNA co-localized with Gag fluctuated (Fig. 2D and 2E). Several studies have described the stochastic nature of HIV-1 transcription with transcriptional bursting, which could explain the variability we observed in the formation of Gag:USvRNA vRNPs^59–62^.

However, it remains unclear why only a small subset of Gag associates with vRNA in the nucleus. One possibility is that the binding of Gag to the USvRNA may act to “mark” the RNA for packaging. It has been demonstrated that packaged genomic RNA adopts different 5’ UTR secondary structures for efficient packaging compared to translated vRNA transcripts depending on the number of guanosines at the 5’ end or stabilization of the polyA stem^63–65^. It would be interesting to investigate whether the binding of Gag to the vRNA in the nucleus promotes certain RNA conformations that encourage packaging. Moreover, previous results showing that HIV-1 Gag localizes to transcriptional burst sites^11^ raise the intriguing question of how co-transcriptional events are coordinated, including recruitment of splicing machinery^66,67^ and the USvRNA export factors Rev and CRM1^68^. It is feasible that Gag association with the USvRNA in the nucleus could act to prevent co-transcriptional splicing from occurring to maintain full-length gRNA by altering the transcriptional profile of cellular genes or recruitment of RNA splicing and nuclear export machinery, tipping the balance toward packaging over subgenomic mRNA export.

Together, the studies presented suggest there may be different populations of Gag; one that remains cytoplasmic and another that undergoes nuclear trafficking to associate with the USvRNA. A recent study exploring Gag-containing complexes (GCCs) indicated that several species of GCCs exist within the cell and bind to various cellular factors with differing contributions to particle formation^69^. Moreover, several mass spectrometry studies have shown Gag association with various host nuclear and cytoplasmic factors^53–57^ (Rice et al., unpublished), supporting the notion that Gag may form transient complexes with different cellular factors for various functions. Of note, the apparent molecular weight of nuclear Gag expressed in J-Lat 10.6 cells appears to be slightly smaller compared to cytoplasmic Gag (Fig. 4A). This slight difference in mobility may be due to a post-translational modification that alter Gag trafficking, localization, or function. It was previously reported that the p6 domain acts as a phosphorylation domain for cyclin-dependent kinases^70^. Further investigation is needed to identify the relevance this finding could have for Gag nuclear trafficking.

HIV-1 integration, latency, and reactivation are influenced by the chromatin state of the infected host cell (as reviewed^71^). In a series of mass spectrometry experiments conducted by several independent research groups, Gag was shown to bind to nuclear factors involved in chromatin remodeling and transcription^53–57^ (Rice et al., unpublished). Moreover, histones were found to be packaged in virus particles isolated from infected cells^72^. These data prompted us to investigate whether Gag preferentially localized to dense heterochromatin or looser, transcriptionally active euchromatin. We found that Gag expressed at low concentrations by either doxycycline-induction or treatment with latency reversal agents, such as prostratin and RMD^25,50,51,73^, preferentially localized to the euchromatin in both HeLa and J-Lat 10.6 reactivated cells (Fig. 3 and 5, respectively). Conversely, higher Gag expression resulted in a near equal ratio of Gag in euchromatin and heterochromatin fractions, suggesting that the amount of Gag in the euchromatin is saturable.

The regulation of chromatin conformation is achieved through the methylation and acetylation of histones. Several markers, including H3K4me3 and H3K27ac, associate with actively transcribing regions, whereas H3K9me3 associates with transcriptionally silent regions (as reviewed^74^). HIV-1 proviral DNA undergoes histone loading, particularly with H3 and H4 histones after nuclear import prior to integration^2^. In the present study, our results indicate the HIV-1 Gag preferentially associates with euchromatin markers H3K4me3 and H3K27ac compared to a heterochromatin marker, supporting the chromatin fractionation results (Fig. 6B). As previously discussed, H3K4me3 histone specifically marks active promoter regions whereas H3K27ac marks active enhancer regions^43–45^. Distal enhancers, such as those marked by H3K27ac, physically interact with promoter regions through forced chromatin looping to promote transcription (as reviewed^75^). Interestingly, Gag exhibited slightly higher associations with H3K27ac marks compared to H3K4me3 (Fig. 6B). It was shown that both 5’ and 3’ LTR regions of integrated provirus are enriched with H3K4me3, and HIV-1 expression is dependent on H3K4 methylation by P-TEFb recruited by Tat^76–78^. Little is known about the relationship between H3K27ac and HIV-1 transcription, but the activation of H3K4me3 enriched genes are interdependent on the presence of H3K27ac^78,79^. Therefore, it is plausible that Gag promotes chromatin loop interactions or utilizes enhancer histone markers to locate the active viral transcription site.

HIV-1 Gag primarily co-localizes with euchromatin marks located near the nuclear periphery (Fig 6C). It is possible that the preferential localization of HIV-1 Gag to histone markers in open chromatin is important for the co-transcriptional selection of gRNA. Our hypothesis is further supported when considering that the integration site of HIV-1 is not random, but occurs with higher frequency in active gene regions near the nuclear periphery^1,3^. A previous report demonstrates that proviral activation of HIV-1 alters chromatin accessibility of neighboring genes^80^. Further experiments, such as ChIP-Seq or ATAC-Seq, would be helpful to pinpoint the specific motifs or DNA elements that play a role in directing Gag trafficking within the nucleus and whether Gag itself alters nearby chromatin structure.

In summary, in these studies investigating the temporospatial properties of nuclear Gag, we demonstrated that a subset of nascent HIV-1 Gag undergoes nuclear localization and traffics to actively transcribing regions of the chromosome, where HIV-1 proviral integration preferentially occurs (Fig. 7). It is possible that these events could facilitate the production of new virus particles following latency reversal by selecting USvRNA for packaging prior to nuclear export. However, the precise mechanism by which this phenomenon occurs remains uncertain and requires further investigation to decipher the exact role(s) of HIV-1 Gag nuclear trafficking following reactivation of proviral gene expression in latently infected cells.

**Figure 7.**
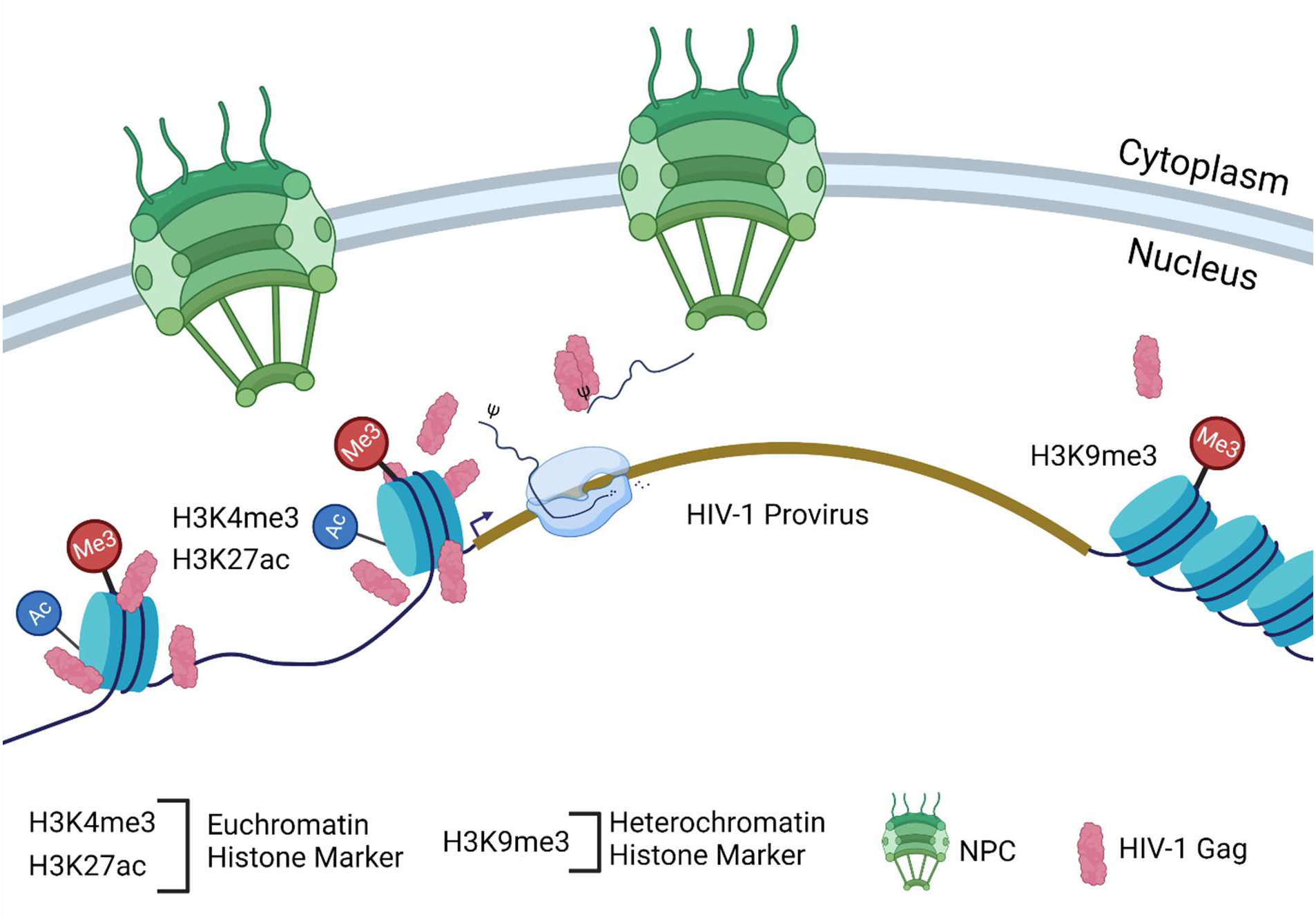
Proposed model of HIV-1 Gag localizing to the nuclear periphery during assembly. Our results demonstrate that HIV-1 Gag is imported shortly after proviral reactivation. Once within the nucleus, HIV-1 Gag preferentially localizes at euchromatin along the nuclear periphery. The model proposes that a subset of HIV-1 Gag produced is imported to the nucleus to the euchromatin where the provirus is most likely integrated. HIV-1 Gag is able co-transcriptionally the USvRNA that will be packaged into new virions.

## Acknowledgements

Research reported in this publication was supported by the National Institute On Drug Abuse of the National Institutes of Health under Award Number R21DA053689 to LJP. The content is solely the responsibility of the authors and does not necessarily represent the official views of the National Institutes of Health. The work was also partially funded by a training grant NCI T32 CA060395-22 to JC (PI, Craig Meyers). We would like to acknowledge those who contributed to this study. All confocal imaging was performed at the Advanced Light Microscopy Core (RRID:SCR_022526), which is funded, in part, by the Pennsylvania State University College of Medicine via the Office of the Vice Dean of Research and Graduate Students and the Pennsylvania Department of Health using Tobacco Settlement Funds (CURE). The content is solely the responsibility of the authors and does not necessarily represent the official views of the University. The Pennsylvania Department of Health specifically disclaims responsibility for any analyses, interpretations, or conclusions. We would like to thank Alan Cochrane and Nicholas Buchkovich for constructive comments on the manuscript, and Parent laboratory members for their important contributions to this work (Rebecca Maldonado and Greg Lambert for technical input, data interpretation, and constructive comments on the manuscript, and Malgorzata Sudol for technical assistance).

## References

1. Dieudonné M, Maiuri P, Biancotto C, et al. Transcriptional competence of the integrated HIV-1 provirus at the nuclear periphery. EMBO Journal. 2009;28(15):2231–2243. doi:10.1038/emboj.2009.141

2. Machida S, Depierre D, Chen HC, et al. Exploring histone loading on HIV DNA reveals a dynamic nucleosome positioning between unintegrated and integrated viral genome. Proc Natl Acad Sci U S A. 2020;117(12):6822–6830. doi:10.1073/pnas.1913754117

3. Albanese A, Arosio D, Terreni M, Cereseto A. HIV-1 pre-integration complexes selectively target decondensed chromatin in the nuclear periphery. PLoS One. 2008;3(6). doi:10.1371/journal.pone.0002413

4. Rabson AB, Graves BJ. Synthesis and Processing of Viral RNA. (Coffin JM, Hughes SH, Varmus HE, eds.). Cold Spring Harbor Laboratory Press; 1997.

5. D’Souza V, Summers MF. How retroviruses select their genomes. Nat Rev Microbiol. 2005;3(8):643–655. doi:10.1038/nrmicro1210

6. Mattei S, Schur FKM, Briggs JAG. Retrovirus maturation - An extraordinary structural transformation. Curr Opin Virol. 2016;18:27–35. doi:10.1016/j.coviro.2016.02.008

7. Jouvenet N, Lainé S, Vivares LP, Mougel M. Cell biology of retroviral RNA packaging. RNA Biol. 2011;8(4):572–580. doi:10.4161/rna.8.4.16030

8. Muriaux D, Darlix JL. Properties and functions of the nucleocapsid protein in virus assembly. RNA Biol. 2010;7(6):744–753. doi:10.4161/rna.7.6.14065

9. Grewe B, Hoffmann B, Ohs I, et al. Cytoplasmic Utilization of Human Immunodeficiency Virus Type 1 Genomic RNA Is Not Dependent on a Nuclear Interaction with Gag. J Virol. 2012;86(6):2990–3002. doi:10.1128/JVI.06874-11

10. Lochmann TL, Bann D v, Ryan EP, et al. NC-mediated nucleolar localization of retroviral gag proteins. Virus Res. 2013;171:304–318.

11. Tuffy KM, Kaddis Maldonado RJ, Chang J, Rosenfeld P, Cochrane A, Parent LJ. HIV-1 gag forms ribonucleoprotein complexes with unspliced viral RNA at transcription sites. Viruses. 2020;12(11):1–26. doi:10.3390/v12111281

12. Scheifele LZ, Garbitt RA, Rhoads JD, Parent LJ. Nuclear entry and CRM1-dependent nuclear export of the Rous sarcoma virus Gag polyprotein. Proceedings of the National Academy of Sciences. 2002;99(6):3944–3949. doi:10.1073/pnas.062652199

13. Scheifele LZ, Kenney SP, Cairns TM, Craven RC, Parent LJ. Overlapping Roles of the Rous Sarcoma Virus Gag p10 Domain in Nuclear Export and Virion Core Morphology. J Virol. 2007;81(19):10718–10728. doi:10.1128/JVI.01061-07

14. Maldonado RJK, Rice B, Chen EC, et al. Visualizing Association of the Retroviral Gag Protein with Unspliced Viral RNA in the Nucleus. mBio. 2020;11(2):1–19. doi:10.1128/mBio.00524-20

15. Garbitt-Hirst R, Kenney SP, Parent LJ. Genetic evidence for a connection between Rous sarcoma virus gag nuclear trafficking and genomic RNA packaging. J Virol. 2009;83(13):6790–6797. doi:10.1128/JVI.00101-09

16. Scheifele LZ, Ryan EP, Parent LJ. Detailed Mapping of the Nuclear Export Signal in the Rous Sarcoma Virus Gag Protein. J Virol. 2005;79(14):8732–8741. doi:10.1128/JVI.79.14.8732

17. Butterfield-Gerson KL, Scheifele LZ, Ryan EP, Hopper AK, Parent LJ. Importin-B Family Members Mediate Alpharetrovirus Gag Nuclear Entry via Interactions with Matrix and Nucleocapsid. J Virol. 2006;80(4):1798–1806. doi:10.1128/JVI.80.4.1798-1806.2006

18. Nash MA, Meyer MK, Decker GL, Arlinghaus RB. A subset of Pr65gag is nucleus associated in murine leukemia virus-infected cells. J Virol. 1993;67(3):1350–1356.

19. Spector DL. Nuclear Domains. J Cell Sci. 2001;114(16):2891–2893.

20. Venkatesh S, Workman JL. Histone exchange, chromatin structure and the regulation of transcription. Nat Rev Mol Cell Biol. 2015;16(3):178–189. doi:10.1038/nrm3941

21. Ghosh RP, Meyer BJ. Spatial Organization of Chromatin: Emergence of Chromatin Structure during Development. Annu Rev Cell Dev Biol. 2021;37:199–232. doi:10.1146/annurev-cellbio-032321-035734

22. Schroder ARW, Shinn P, Chen H, Berry C, Ecker JR, Bushman F. HIV-1 Integration in the Human Genome Favors Active Genes and Local Hotspots. Cell. 2002;110(4):521–529.

23. Verdin E, Paras P, Van Lint C. Chromatin Disruption in the Promoter of HIV-1 During Transcriptional Activation. EMBO J. 1993;12(8):3249–3259.

24. Van Lint C, Emiliani S, Ott M, Verdin E. Transcriptional activation and chromatin remodeling of the HIV-I promoter in response to histone acetylation. EMBO J. 1997;15(5):1112–1120.

25. Van Lint C, Emiliani S, Verdin E. The expression of a small fraction of cellular genes is changed in response to histone hyperacetylation. Gene Expr. 1996;5(4-5):245–253.

26. Taube R, Peterlin BM. Lost in transcription: Molecular mechanisms that control HIV latency. Viruses. 2013;5(3):902–927. doi:10.3390/v5030902

27. Elliott JH, Wightman F, Solomon A, et al. Activation of HIV Transcription with Short-Course Vorinostat in HIV-Infected Patients on Suppressive Antiretroviral Therapy. PLoS Pathog. 2014;10(11). doi:10.1371/journal.ppat.1004473

28. Archin NM, Liberty AL, Kashuba AD, et al. Administration of vorinostat disrupts HIV-1 latency in patients on antiretroviral therapy. Nature. 2012;487(7408):482–485. doi:10.1038/nature11286

29. Archin NM, Kirchherr JL, Sung JAM, et al. Interval dosing with the HDAC inhibitor vorinostat effectively reverses HIV latency. Journal of Clinical Investigation. 2017;127(8):3126–3135. doi:10.1172/JCI92684

30. Søgaard OS, Graversen ME, Leth S, et al. The Depsipeptide Romidepsin Reverses HIV-1 Latency In Vivo. PLoS Pathog. 2015;11(9):1–22. doi:10.1371/journal.ppat.1005142

31. Col E, Caron C, Seigneurin-Berny D, Gracia J, Favier A, Khochbin S. The Histone Acetyltransferase, hGCN5, Interacts with and Acetylates the HIV Transactivator, Tat. Journal of Biological Chemistry. 2001;276(30):28179–28184. doi:10.1074/jbc.M101385200

32. Hottiger MO, Nabel GJ. Interaction of Human Immunodeficiency Virus Type 1 Tat with the Transcriptional Coactivators p300 and CREB Binding Protein. J Virol. 1998;72(10):8252–8256. doi:10.1128/jvi.72.10.8252-8256.1998

33. Benkirane M, Chun RF, Xiao H, et al. Activation of integrated provirus requires histone acetyltransferase: p300 and P/CAF are coactivators for HIV-1 Tat. Journal of Biological Chemistry. 1998;273(38):24898–24905. doi:10.1074/jbc.273.38.24898

34. Lachner M, O’Carroll D, Rea S, Karl M, Jenuwein T. Methylation of histone H3 lysine 9 creates a binding site for HP1 proteins. Letters to Nature. 2001;410:116–120.

35. Marban C, Redel L, Suzanne S, et al. COUP-TF interacting protein 2 represses the initial phase of HIV-1 gene transcription in human microglial cells. Nucleic Acids Res. 2005;33(7):2318–2331. doi:10.1093/nar/gki529

36. Azzaz AM, Vitalini MW, Thomas AS, et al. Human heterochromatin protein 1α promotes nucleosome associations that drive chromatin condensation. Journal of Biological Chemistry. 2014;289(10):6850–6861. doi:10.1074/jbc.M113.512137

37. Rohr O, Lecestre D, Chasserot-Golaz S, et al. Recruitment of Tat to Heterochromatin Protein HP1 via Interaction with CTIP2 Inhibits Human Immunodeficiency Virus Type 1 Replication in Microglial Cells. J Virol. 2003;77(9):5415–5427. doi:10.1128/jvi.77.9.5415-5427.2003

38. Friedman J, Cho WK, Chu CK, et al. Epigenetic Silencing of HIV-1 by the Histone H3 Lysine 27 Methyltransferase Enhancer of Zeste 2. J Virol. 2011;85(17):9078–9089. doi:10.1128/jvi.00836-11

39. Pearson R, Kim YK, Hokello J, et al. Epigenetic Silencing of Human Immunodeficiency Virus (HIV) Transcription by Formation of Restrictive Chromatin Structures at the Viral Long Terminal Repeat Drives the Progressive Entry of HIV into Latency. J Virol. 2008;82(24):12291–12303. doi:10.1128/jvi.01383-08

40. Tripathy MK, McManamy MEM, Burch BD, Archin NM, Margolis DM. H3K27 Demethylation at the Proviral Promoter Sensitizes Latent HIV to the Effects of Vorinostat in Ex Vivo Cultures of Resting CD4 + T Cells. J Virol. 2015;89(16):8392–8405. doi:10.1128/jvi.00572-15

41. Pocock GM, Becker JT, Swanson CM, Ahlquist P, Sherer NM. HIV-1 and M-PMV RNA Nuclear Export Elements Program Viral Genomes for Distinct Cytoplasmic Trafficking Behaviors. PLoS Pathog. 2016;12(4). doi:10.1371/journal.ppat.1005565

42. Chase GP, Rameix-Welti MA, Zvirbliene A, et al. Influenza virus ribonucleoprotein complexes gain preferential access to cellular export machinery through chromatin targeting. PLoS Pathog. 2011;7(9). doi:10.1371/journal.ppat.1002187

43. Heintzman ND, Stuart RK, Hon G, et al. Distinct and predictive chromatin signatures of transcriptional promoters and enhancers in the human genome. Nat Genet. 2007;39(3):311–318. doi:10.1038/ng1966

44. Heintzman ND, Hon GC, Hawkins RD, et al. Histone modifications at human enhancers reflect global cell-type-specific gene expression. Nature. 2009;459(7243):108–112. doi:10.1038/nature07829

45. Creyghton MP, Cheng AW, Welstead GG, et al. Histone H3K27ac separates active from poised enhancers and predicts developmental state. Proc Natl Acad Sci U S A. 2010;107(50):21931–21936. doi:10.1073/pnas.1016071107

46. Williams SA, Chen LF, Kwon H, et al. Prostratin antagonizes HIV latency by activating NF-κB. Journal of Biological Chemistry. 2004;279(40):42008–42017. doi:10.1074/jbc.M402124200

47. Osborn L, Kunkel S, Nabel GJ. Tumor necrosis factor a and interleukin 1 stimulate the human immunodeficiency virus enhancer by activation of the nuclear factor kB. Proc Natl Acad Sci USA. 1989;86:2336–2340.

48. Duh EJ, Maury WJ, Folks TM, Fauci AS, Rabson AB. Tumor Necrosis Factor a Activates Human Immunodeficiency Virus Type 1 through Induction of Nuclear Factor Binding to the NF-KB Sites in the Long Terminal Repeat (ACH2 T-Celi Line/Provirus/Retroviral Transcription). Vol 86.; 1989.

49. Griffen GE, Leung K, Folks TM, Kunkel S, Nabel GJ. Activation of HIV gene expression during monocyte differentiation by induction of NF-kB. Letters to Nature. 1989;339:70–73.

50. Wei DG, Chiang V, Fyne E, et al. Histone Deacetylase Inhibitor Romidepsin Induces HIV Expression in CD4 T Cells from Patients on Suppressive Antiretroviral Therapy at Concentrations Achieved by Clinical Dosing. PLoS Pathog. 2014;10(4). doi:10.1371/journal.ppat.1004071

51. Jønsson KL, Tolstrup M, Vad-Nielsen J, et al. Histone deacetylase inhibitor romidepsin inhibits De Novo HIV-1 infections. Antimicrob Agents Chemother. 2015;59(7):3984–3994. doi:10.1128/AAC.00574-15

52. Kiernan RE, Vanhulle C, Schiltz L, et al. HIV-1 Tat transcriptional activity is regulated by acetylation. EMBO Journal. 1999;18(21):6106–6118. doi:10.1093/emboj/18.21.6106

53. Engeland CE, Brown NP, Börner K, et al. Proteome analysis of the HIV-1 Gag interactome. Virology. 2014;460-461(1):194–206. doi:10.1016/j.virol.2014.04.038

54. Jäger S, Cimermancic P, Gulbahce N, et al. Global landscape of HIV-human protein complexes. Nature. 2012;481(7381):365–370. doi:10.1038/nature10719

55. Ritchie C, Cylinder I, Platt EJ, Barklis E. Analysis of HIV-1 Gag Protein Interactions via Biotin Ligase Tagging. J Virol. 2015;89(7):3988–4001. doi:10.1128/jvi.03584-14

56. Li Y, Frederick KM, Haverland NA, Ciborowski P, Belshan M. Investigation of the HIV-1 matrix interactome during virus replication. Proteomics Clin Appl. 2016;10(2):156–163. doi:10.1002/prca.201400189

57. le Sage V, Cinti A, Valiente-Echeverría F, Mouland AJ. Proteomic analysis of HIV-1 Gag interacting partners using proximity-dependent biotinylation. Virol J. 2015;12(1). doi:10.1186/s12985-015-0365-6

58. Mohammadi P, Desfarges S, Bartha I, et al. 24 Hours in the Life of HIV-1 in a T Cell Line. PLoS Pathog. 2013;9(1). doi:10.1371/journal.ppat.1003161

59. Tantale K, Garcia-Oliver E, Robert MC, et al. Stochastic pausing at latent HIV-1 promoters generates transcriptional bursting. Nat Commun. 2021;12(1). doi:10.1038/s41467-021-24462-5

60. Skupsky R, Burnett JC, Foley JE, Schaffer D v., Arkin AP. HIV promoter integration site primarily modulates transcriptional burst size rather than frequency. PLoS Comput Biol. 2010;6(9). doi:10.1371/journal.pcbi.1000952

61. Burnett JC, Miller-Jensen K, Shah PS, Arkin AP, Schaffer D v. Control of stochastic gene expression by host factors at the HIV promoter. PLoS Pathog. 2009;5(1). doi:10.1371/journal.ppat.1000260

62. Singh A, Razooky B, Cox CD, Simpson ML, Weinberger LS. Transcriptional bursting from the HIV-1 promoter is a significant source of stochastic noise in HIV-1 gene expression. Biophys J. 2010;98(8). doi:10.1016/j.bpj.2010.03.001

63. Nikolaitchik OA, Liu S, Kitzrow JP, et al. Selective packaging of HIV-1 RNA genome is guided by the stability of 5 0 untranslated region polyA stem. doi:10.1073/pnas.2114494118/-/DCSupplemental

64. Nikolaitchik OA, Somoulay X, Rawson JMO, Yoo JA, Pathak VK, Hu WS. Unpaired Guanosines in the 5′ Untranslated Region of HIV-1 RNA Act Synergistically To Mediate Genome Packaging. J Virol. 2020;94(21). doi:10.1128/jvi.00439-20

65. Brown JD, Kharytonchyk S, Chaudry I, et al. Structural Basis for Transcriptional Start Site Control of HIV-1 RNA Fate. https://www.science.org

66. Ameur A, Zaghlool A, Halvardson J, et al. Total RNA sequencing reveals nascent transcription and widespread co-transcriptional splicing in the human brain. Nat Struct Mol Biol. 2011;18(12):1435–1440. doi:10.1038/nsmb.2143

67. Tilgner H, Knowles DG, Johnson R, et al. Deep sequencing of subcellular RNA fractions shows splicing to be predominantly co-transcriptional in the human genome but inefficient for lncRNAs. Genome Res. 2012;22(9):1616–1625. doi:10.1101/gr.134445.111

68. Nawroth I, Mueller F, Basyuk E, et al. Stable assembly of HIV-1 export complexes occurs cotranscriptionally. RNA. 2014;20(1):1–8. doi:10.1261/rna.038182.113

69. Deng Y, Hammond JA, Pauszek R, et al. Discrimination between Functional and Non-functional Cellular Gag Complexes involved in HIV-1 Assembly: Characterizing Intracellular HIV-1 Gag Complexes. J Mol Biol. 2021;433(8). doi:10.1016/j.jmb.2021.166842

70. Müller B, Patschinsky T, Kräusslich HG. The Late-Domain-Containing Protein p6 Is the Predominant Phosphoprotein of Human Immunodeficiency Virus Type 1 Particles. J Virol. 2002;76(3):1015–1024. doi:10.1128/jvi.76.3.1015-1024.2002

71. Agosto LM, Gagne M, Henderson AJ. Impact of chromatin on HIV replication. Genes (Basel). 2015;6(4):957–976. doi:10.3390/genes6040957

72. Chertova E, Chertov O, Coren L v., et al. Proteomic and Biochemical Analysis of Purified Human Immunodeficiency Virus Type 1 Produced from Infected Monocyte-Derived Macrophages. J Virol. 2006;80(18):9039–9052. doi:10.1128/jvi.01013-06

73. Reuse S, Calao M, Kabeya K, et al. Synergistic activation of HIV-1 expression by deacetylase inhibitors and prostratin: Implications for treatment of latent infection. PLoS One. 2009;4(6). doi:10.1371/journal.pone.0006093

74. Bannister AJ, Kouzarides T. Regulation of chromatin by histone modifications. Cell Res. 2011;21(3):381–395. doi:10.1038/cr.2011.22

75. Schoenfelder S, Fraser P. Long-range enhancer–promoter contacts in gene expression control. Nat Rev Genet. 2019;20(8):437–455. doi:10.1038/s41576-019-0128-0

76. Park J, Lim CH, Ham S, Kim SS, Choi BS, Roh TY. Genome-wide analysis of histone modifications in latently HIV-1 infected T cells. AIDS. 2014;28(12):1719–1728. doi:10.1097/QAD.0000000000000309

77. Zhou M, Deng L, Lacoste V, et al. Coordination of Transcription Factor Phosphorylation and Histone Methylation by the P-TEFb Kinase during Human Immunodeficiency Virus Type 1 Transcription. J Virol. 2004;78(24):13522–13533. doi:10.1128/jvi.78.24.13522-13533.2004

78. Zhao W, Xu Y, Wang Y, et al. Investigating crosstalk between H3K27 acetylation and H3K4 trimethylation in CRISPR/dCas-based epigenome editing and gene activation. Sci Rep. 2021;11(1). doi:10.1038/s41598-021-95398-5

79. Tie F, Banerjee R, Stratton CA, et al. CBP-mediated acetylation of histone H3 lysine 27 antagonizes Drosophila Polycomb silencing. Development. 2009;136(18):3131–3141. doi:10.1242/dev.037127

80. Shah R, Gallardo CM, Jung YH, et al. Activation of HIV-1 proviruses increases downstream chromatin accessibility. iScience. 2022;25(12). doi:10.1016/j.isci.2022.105490

